# Modifying Pavlovian-To-Instrumental Transfer By Approach Avoidance Training In Healthy Subjects – A Proof of Concept Study

**DOI:** 10.1101/2022.07.01.497725

**Authors:** Annika Rosenthal, Ke Chen, Anne Beck, Nina Romanczuk-Seiferth

## Abstract

The modulation of instrumental action by conditioned Pavlovian cues is hypothesized to play a role in the emergence and maintenance of maladaptive behavior. The Pavlovian-to-instrumental transfer (PIT) task is designed to examine the magnitude of the influence of cues on behavior and we aim to manipulate the motivational value of Pavlovian cues to reduce their effect on instrumental responding. To this end, we utilized a joystick-based modification of approach and avoidance propensities that has shown success in clinical populations. In 35 healthy subjects, we examined changes in PIT after completion of either avoidance training or sham training. We found no effect of training on approach avoidance propensities but higher response rate towards negative stimuli during PIT after systematic avoidance training compared to sham training. On the other hand, we saw an increased PIT effect after sham training. These results imply that training can alter the strength of the influence of cues on instrumental behavior and suggest training to be beneficial in reducing environmental triggers of maladaptive behavior.

## Introduction

Stimuli that are associated with rewards have been demonstrated to encourage behaviors attributed to past rewarding experiences ^1,2^. This so-called concept of cue-reactivity is central to human and animal adaptive behavior such as food seeking or reproduction ^2^. However, cue-reactivity has been mostly researched in the background of disparaging behavior such as substance abuse, binge eating or other behavioral patterns marked by conflicts of behavioral goals and value assigned to stimuli ^3-5^. For instance, after initially producing rewarding effects, prolonged drug abuse could alter motivational drive and sensitize to drug-related conditioned responding and craving ^6,7^. These factors play a fundamental role in the development and maintenance of substance use disorders (SUDs). The interplay of instrumental behavior and reinforcing properties of stimuli has been conceptualized in various ways. One way cue-motivated behavior has been modeled, is through Pavlovian-to-instrumental transfer (PIT) ^8,9^. Essentially, PIT tasks constitute of an instrumental training to establish response–outcome (R–O) associations by linking responses to reward delivery. In a Pavlovian conditioning phase, previously neutral stimuli (CS) are coupled to rewarding outcomes (US), to establish stimulus–outcome (CS–US) associations. In the transfer test, instrumental behavior is assessed in extinction and in the presence of the outcome-associated stimuli ^10^. An increase of instrumental behavior due to high motivational salience of the Pavlovian CS has been found in various clinical populations as well as at-risk groups, i.e. alcohol use disorder ^11^, social drinking ^12^ and obesity ^13^. Aversive PIT was exaggerated in patients with depression ^14^ and increased loss aversion PIT was found in subjects with gambling disorder ^15^. However, the underlying mechanisms of the nature of transfer effects are debated ^16^. Associative accounts postulate the dissociative engagement of motivational and cognitive control and within this context, biased action selection towards reward cues has been explained ^17^. Similarly, originating within a dual-systems framework, cognitive bias modification (CBM) was developed with the idea to evaluate these biases in the context of maladaptive behavior ^18^. One form of CBM focuses on bias in the automatically activated action tendency to approach or avoid certain stimuli. This approach avoidance task (AAT) operationalizes push (avoid) and pull (approach) behavior with a joystick experiment in which subjects are instructed to react to content or content-unrelated features of a stimulus. Studies have shown that an increased approach bias towards drug-related stimuli was related to drug consumption or addiction severity ^19,20^ as well as food-associated stimuli increased approach bias in food craving ^21^ and was correlated to uncontrolled eating ^22^. In this context, modified versions of the AAT have been used to successfully retrain these increased action tendencies in clinical populations ^23-25^. An indication of efficacy is uncertain, however, due to mixed results for reviews, see ^26,27^. In extension of this, Pavlovian conditioning of previously neutral cues has been shown to elicit significant approach tendencies towards these cues as well. For example, approach bias was enhanced when participants were faced with abstract stimuli that were previously paired with chocolate ^28^ or tobacco ^29^.

Regarding the role of implicit motivational processes that drive behavior, PIT effects and approach bias have been theorized to influence each other but the exact nature of this has not been disentangled, yet. While it has been proposed that approach bias plays a role in PIT ^30^, it has also been stipulated that transfer effects drive approach bias ^31^. Considering clinical relevance, increased PIT effects in psychiatric populations, especially in the addiction domain, have not been subjected to systematic modification, yet. It has been proposed, however, that clinical populations with strong PIT effects could profit from approach modification training to reduce PIT effects and increase behavioral control ^11^. From a therapeutic perspective, it makes sense to systematically reduce the effect of Pavlovian cues that trigger maladaptive behavior: On the one hand, to prevent possible situational habit formation in at-risk populations and on the other hand, to disrupt the effect of environmental cues and ensure abstinence from already established dysfunctional behavior.

In light of this, we want to investigate whether PIT effects can be translated into approach and avoidance biases towards previously conditioned cues in the context of an AAT and subsequent retraining of these biases. In order to examine this in a healthy population, we first want to evaluate whether Pavlovian conditioning of neutral cues in the context of a PIT task elicits approach and avoidance biases toward them. In a next step, we want to find out whether these action tendencies can be manipulated via a modified training version of the AAT. Lastly, we want to assess the effects of this systematic manipulation of the approach avoidance propensities of Pavlovian cues on PIT processes in healthy adults.

## Methods

All procedures complied with the Declaration of Helsinki and were approved by the ethical committee of the Charité - Universitätsmedizin Berlin. All participants gave fully written informed consent.

### Participants

The study was conducted in Berlin, Germany, and all participants were recruited through internet advertisement. Participants were included between 18 and 65 years of age and if they were neither pregnant nor breastfeeding. Exclusion criteria were current or past SCID (Structured Clinical Interview for DSM) disorders, including personality disorders, and SUD (except tobacco use disorder (TUD) and mild (up to 6 criteria) cannabis use disorder (CUD)). The sample consisted of 35 participants (22 female) and age ranged from 19 to 60 (mean = 35,9 SD = 12,01) (*see table 1*.).

**Table 1.**
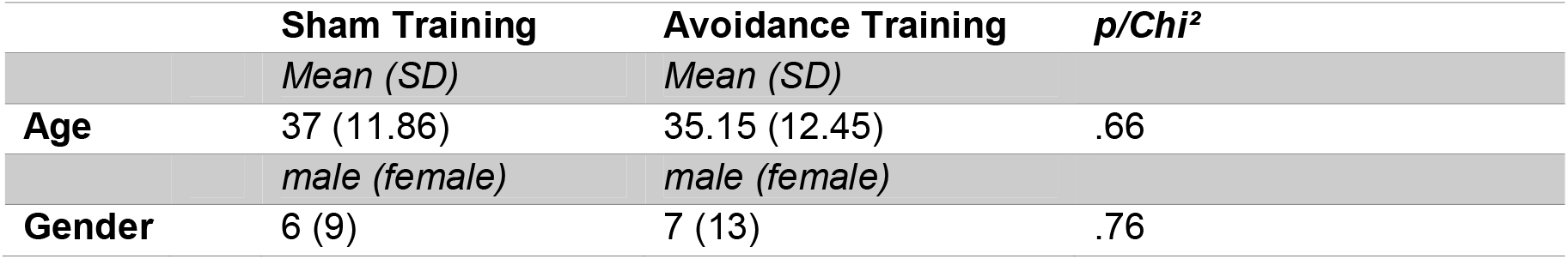
Descriptives of sham- and avoidance training group

|  | **Sham Training** | **Avoidance Training** | ***p/Chi<sup>2</sup>*** |
| --- | --- | --- | --- |
|  | *Mean (SD)* | *Mean (SD)* |  |
| **Age** | 37 (11.86) | 35.15 (12.45) | .66 |
|  | *male (female)* | *male (female)* |  |
| **Gender** | 6 (9) | 7 (13) | .76 |

### Procedure

The PIT paradigm was administered in six parts, which consisted of: (1) instrumental training, (2) Pavlovian training, (3, 5) PIT before and after AAT training and (4, 6) a forced choice task before and after AAT training (*see* *Fig*.□*<u>1</u>A-F*). The task was programmed with Matlab 2019b (MATLAB version 9.7.0, 2019; MathWorks, Natick, MA, USA) using the Psychophysics Toolbox Version 3 (PTB 3.0.15) extension ^32-34^. For an extensive description of the PIT paradigm, please see Garbusow *et*□*al*· ^35^. After the Pavlovian conditioning phase (*1B*) we administered the AAT (*2A*) to assess the participants’ approach/avoidance bias. Following PIT pretest (*1C*) and the forced choice task (query trials), the AAT training and post-test (*2B*) were administered.

**Figure 1.**
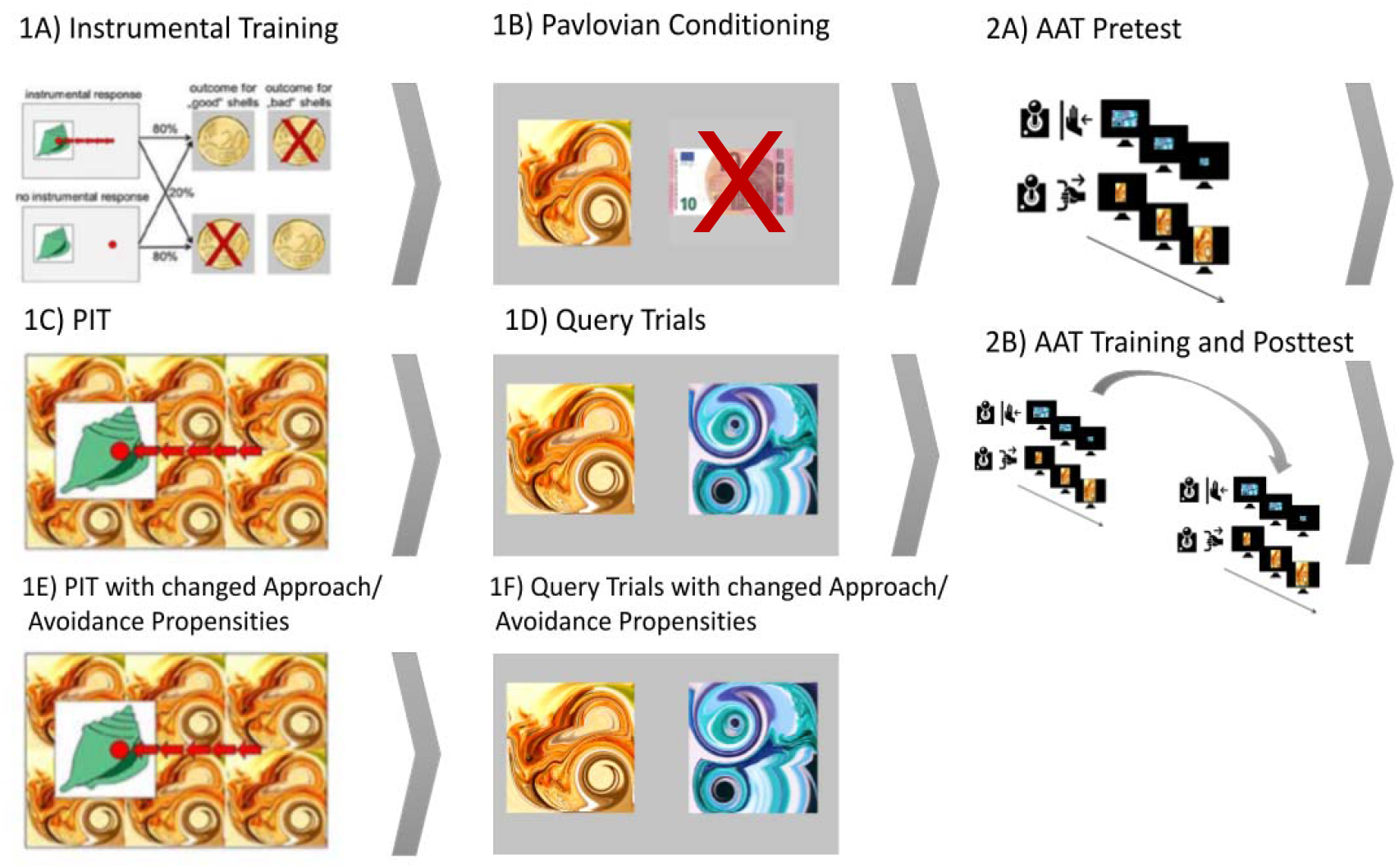
Schematic representation of the study design. Participants first conducted the instrumental training (1A). Pavlovian conditioning (1B) is followed by an AAT pretest (2A). After PIT and Query trials (1C,1D), AAT training and posttest (2B) is followed by a second PIT and Query trials (1E,1F). The grey arrows indicate the temporal order of the tasks.

**Figure 2.**
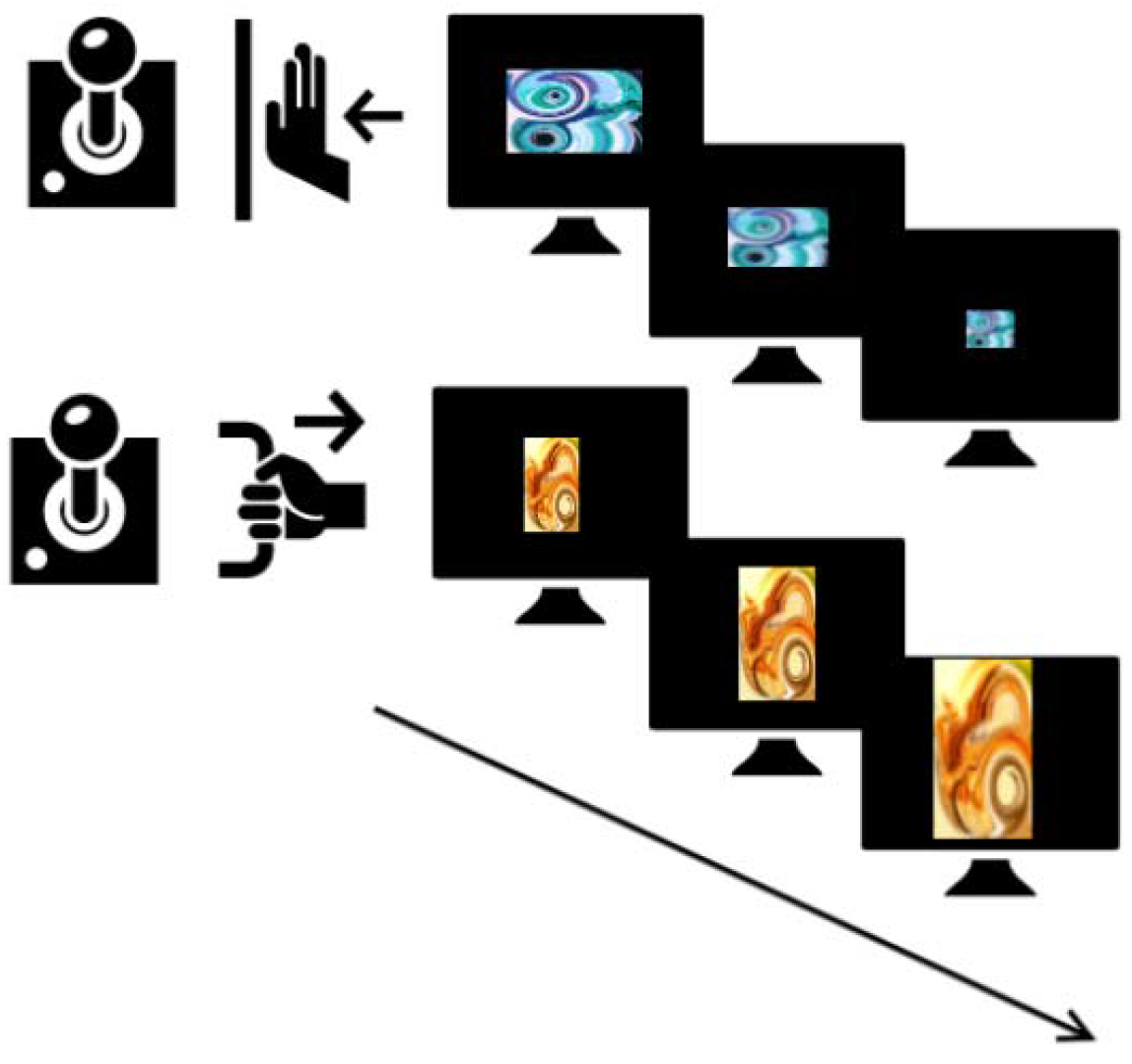
In detail representation of the AAT training, i.e. the avoidance training. Positively conditioned fractals are pushed away (top) and negatively conditioned fractals are pulled towards the subject (bottom).

### Instrumental training

Instrumental stimuli consisted of shells in various colors and shapes that were presented on a computer screen. The instructions to the participants were to collect “good” and leave “bad” shells while receiving probabilistic feedback. In order to collect a good shell, the subjects had to repeatedly press the left mouse button (at least 5 times), while they had to omit a reaction when a bad shell was presented (0-4 button presses were counted as omission). In random fashion, correct responses were rewarded with 20 Cents in 80% of the trials and punished with a loss of 20 Cents in 20% of trials, and vice versa for incorrect responses. Dependent on performance, a learning criterion (80% correct trials over 16 consecutive trials) determined the task length to be between 60 and 120 trials.

### Pavlovian Conditioning

Subjects were presented with 48 trials of abstract image-sound combinations (compound CS) that were paired with monetary reward or punishment (US). The compound CS was presented for 3 seconds on the left or right side of the screen, followed by a delay of 3 seconds with two fixation crosses at the two potential CS locations, then a US (monetary reward 10€, punishment -10€ or no stimulus) was presented for another 3 seconds. Subjects had to passively watch and memorize the CS and US pairings. Abstract pictures and associated US were randomly paired.

### Pavlovian-to-instrumental transfer

The instrumental task was performed in formal extinction, which means that the monetary outcome was not shown. During the trials the background was alternately tiled with one of the CS. The task had a duration of 108 trials with each lasting 3 seconds.

### Query trials

After the transfer stage, a forced choice task was administered to confirm the success of the Pavlovian conditioning. Based on their subjective preference, subjects had to choose between two Pavlovian CSs (9 trials). All pairings were presented in an interleaved, randomized order.

### AAT

This version of the AAT was programmed so that participants had to respond according to the orientation of the pictures (*see Fig. 3*). With a joystick, all horizontal pictures were to be pushed away (avoidance), and all vertical pictures were to be pulled closer (approach). To mimic realistic approach/avoidance behavior, a zoom feature was used. In the approach (pull) movement, pictures grew larger; while pushing resulted in shrinking pictures. The task structure was adapted from Wiers et al. ^36^. As stimuli, we used the same abstract pictures (CS) as in the PIT task. The AAT consisted of three different phases: a pre-test AAT, a training AAT and a post-test AAT. A practice phase of 40 trials, in which participants learned to move the joystick according to orientation, was followed by 120 trials in which all three abstract pictures had to be pushed/pulled in equal frequency. During the training phase that consisted of 300 trials, participants moved the joystick according to the assigned condition. In the sham condition, stimuli were shown equally often in all possible trial types (push or pull). In the avoidance condition, participants had to push positively conditioned pictures (10€) and pull negatively conditioned pictures (−10€) while neutral pictures were both pushed and pulled in equal frequency. In the post-test phase, 120 trials with equal frequency of all possible trial types were administered.

## Analysis

### Statistical Analysis

Data were analyzed in Matlab 2011a and Jamovi (The Jamovi Project, 2022). Generalized linear modeling (GLM) as well as generalized linear mixed modeling (GLMM) was used since the reaction time (RT) data from the AAT were non-normally distributed and the PIT response showed a zero inflation as no response was required on some of the trials. All models used log link functions.

#### AAT

In order to exploratorily assess the individual contribution of Pavlovian values on approach avoidance behavior at pre and post training separately, initial gamma distributed models were applied to test for the effect of background CS (−10€/neutral/+10€; coded as −0.5, 0, +0.5), direction (push/pull; coded as -0.5/+0.5) as well as their interaction on RT in each trial. Subsequently a GLMM, a gamma distributed model was built in which the RT in each trial was predicted by the value of the background CS (−10€/neutral/+10€; coded as −0.5, 0, +0.5) the direction (push/pull; coded as -0.5/+0.5), the training condition (Sham/Avoidance; coded as -0.5/+0.5) and time (pre Training/post Training; coded as -0.5/+0.5) as well as their interactions. Initial model comparison based on Akaike Information Criterion (AIC) ^37^ of different random-effects structures indicated best model fit for taking intercept, main effects of CS value, direction and time as random effects across subjects.

#### PIT

Initial model fitting indicated a Poisson distributed model to be overdispersed (*Chi*^*2*^*/DF* = 3.77) ^38^. To account for this, a model with negative binomial distribution was built to assess the individual contribution of Pavlovian values on behavior. Here, the number of button presses in each trial was predicted by the value of the background CS (−10€/neutral/+10€; coded as −0.5, 0, +0.5) the instrumental condition (not collect/collect; coded as -0.5/+0.5), the training condition (Sham/Avoidance; coded as -0.5/+0.5) and time (pre Training/post Training; coded as -0.5/+0.5) as well as their interactions. Model comparisons indicated the within-subject factors intercept, main effect of CS value and time to be the best fitting random factor structure.

## Results

### AAT

Of the experimental trials, 6,92% had to be excluded (first and last percentile as well as all participants with commission errors > 21). These exclusion criteria are in line with previous studies that investigated AAT data ^25,39^.

We did not find a significant effect of the background CS, direction or their interaction on RT at pretest (*see table 2*). However, descriptively, in the pull direction, participants seemed to approach (lower RT) positive CS faster (mean = 682.949, SE = 6.551) in comparison to neutral (mean = 689.956, SE = 6.665) and negative CS (mean = 699.248, SE = 6.744)

**Table 2.**
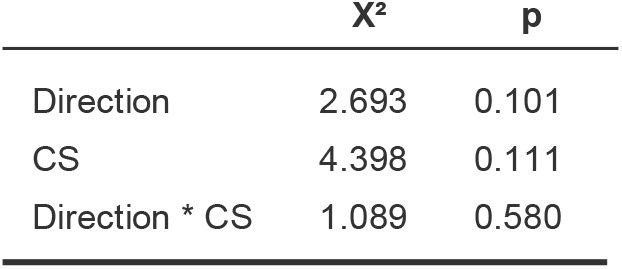
Results of generalized linear model testing the interaction effect of response direction and conditioned stimulus on reaction time during the AAT pretest.

In our GLMM, the main effect of interest was the interaction between training by time by CS by direction. This however, was not significant, indicating there was no effect of training on the change of approach or avoidance bias towards the CS (*please see table 3 and figure 3*). However, there was a main effect of time (*estimate* = -0.07456; SE = 0.02503; *p* = □.003), indicating a general decrease of RT from pre AAT to post AAT. Additionally, now the direction by CS interaction was significant (*X*^*2*^ *=* 7.5775; p = 0.023), here the RT in the avoidance direction is trend wisely higher for the negative CS than neutral CS (*estimate* = - 0.01647; SE = 0.00967; *p* = □.089) but lower for the negative CS compared to the neutral CS in the approach direction, however, not significant (*estimate* = 0.0810; SE = 0.00976; *p* = □.407). Additionally the CS by direction by time interaction was significant (*X*^*2*^ = 6.1437; *p* = .046), indicating that irrespective of training, the RT of avoidance of negative CS was significantly lower compared to avoidance RT towards positive CS (*estimate* = -0.02825; SE = 0.0138; *p* = □.041).

**Table 3.** Results of generalized linear mixed model testing the interaction effect between response direction, conditioned stimulus, time and training on reaction time in the AAT.

| | $X^2$ | $p$ |
| --- | --- | --- |
| Direction | 2.0985 | 0.147 |
| CS | 0.5692 | 0.752 |
| time | 8.8714 | **0.003** |
| Training | 0.0489 | 0.825 |
| Direction * CS | 7.5775 | **0.023** |
| Direction * time | 0.0337 | 0.854 |
| CS * time | 2.5172 | 0.284 |
| Direction * Training | 0.4277 | 0.513 |
| CS * Training | 0.4546 | 0.797 |
| time * Training | 0.0731 | 0.787 |
| Direction * CS * time | 6.1437 | **0.046** |
| Direction * CS * Training | 1.0985 | 0.577 |
| Direction * time * Training | 1.9854 | 0.159 |
| CS * time * Training | 0.6203 | 0.733 |
| Direction * CS * time * Training | 2.6757 | 0.262 |

**Figure 2.**
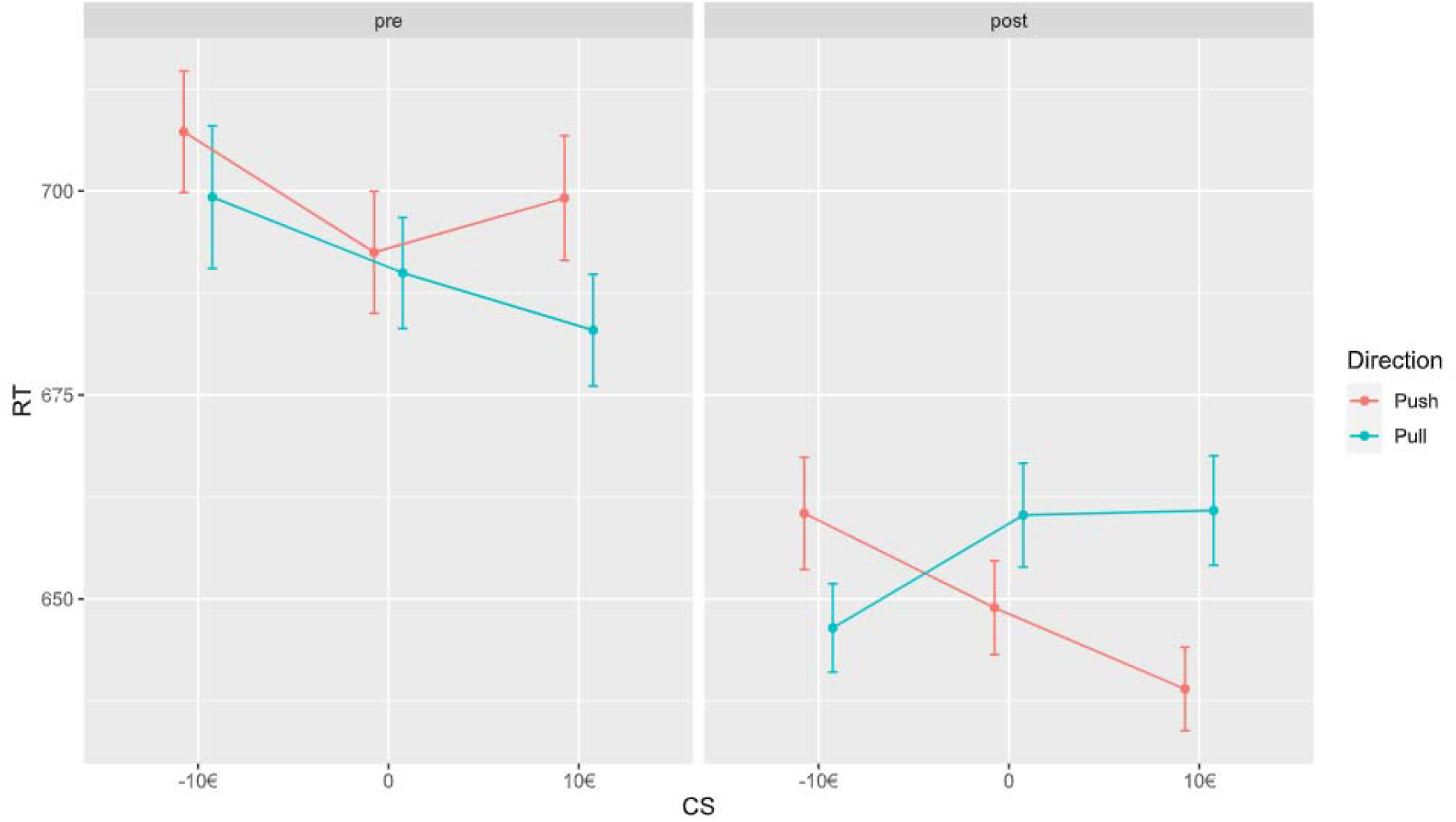
There was an overall effect of time, participants responded faster in the post AAT. In addition, after training, there was an increase in RT towards positive compared to negative CS.

Please see Fig.3 for a graphic representation of the results of this interaction.

**Figure 3.**
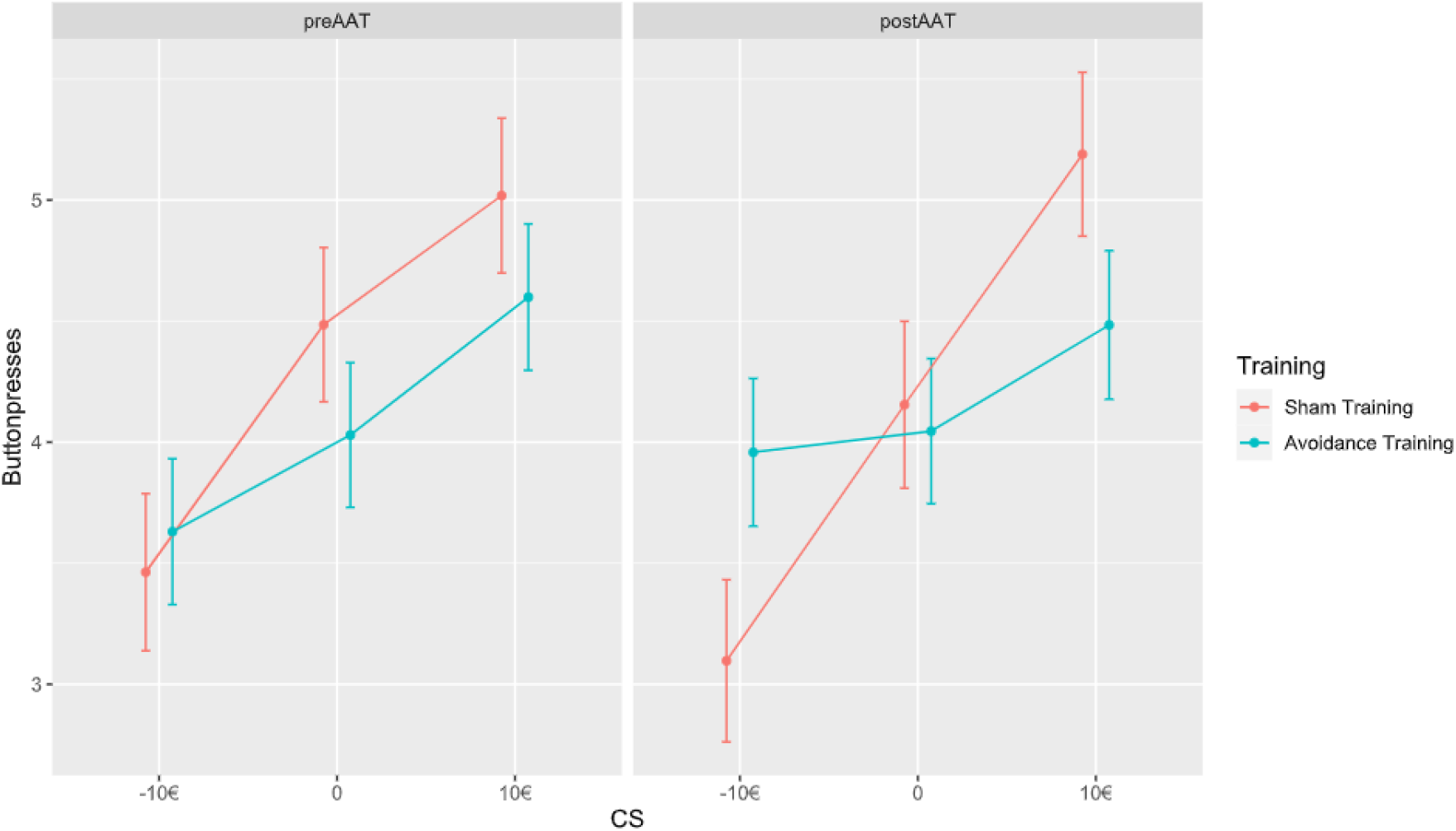
Graph depicting the PIT effect before and after AAT sham/avoidance training. There was a significant interaction effect of CS by time by training, which indicates that the instrumental behavior changed over time due to training.

We explored the effects in GLM with post AAT data only and found no significant interaction between CS by direction by training. However, in an exploratory fashion, we examined simple effects and found avoidance training to decrease the RT of in push trials when positive CS were displayed (estimate =-0.049; *z* = -2.90; p = .004) (*supplementary Fig*.*1*).

#### PIT

We found a significant effect of the background CS and instrumental condition but no interaction on the number of button presses at PIT pretest (*table 4*). As expected, subjects responded with more button presses in positive CS trials (mean = 4.30; SE = 0.154) than in neutral CS trials (mean = 3.68; SE = 0.134) and negative CS trials (mean = 3.05; SE = 0.114). Button presses in collect trials were higher (mean = 6.19; SE = 0.175) than in not collect trials (mean = 2.14; SE = 0.670).

**Table 4.** Results of the pre training PIT.

| | $\chi^2$ | p |
| --- | --- | --- |
| CS | 43.53 | <.001 |
| Instrumental Condition | 626.20 | <.001 |
| CS * Instrumental Condition | 2.97 | 0.226 |

**Table 5.** Results of generalized linear mixed model testing the interaction effect of instrumental response, conditioned stimulus, time and training on button presses in the PIT.

|  | X<sup>2</sup> | p |
| --- | --- | --- |
| CS | 11.2733 | **0.004** |
| Time | 4.2642 | **0.039** |
| Training | 2.0776 | 0.149 |
| Instrumental Condition | 552.2536 | **<.001** |
| CS * time | 3.9257 | 0.140 |
| CS * Training | 3.7589 | 0.153 |
| time * Training | 11.2894 | **<.001** |
| CS * Instrumental Condition | 4.1250 | 0.127 |
| time * Instrumental Condition | 6.9765 | **0.008** |
| Training * Instrumental Condition | 0.0201 | 0.887 |
| CS * time * Training | 10.1443 | **0.006** |
| CS * time * Instrumental Condition | 4.8515 | 0.088 |
| CS * Training * Instrumental Condition | 0.3972 | 0.820 |
| time * Training * Instrumental Condition | 3.7041 | 0.054 |
| CS * time * Training * Instrumental Condition | 1.5239 | 0.467 |

The GLMM focused on the training effect on PIT showed significant main effects of CS (*x*^*2*^ =11.273; *p* = .004), time (*x*^*2*^ =4.264; *p* = .039) and instrumental condition (*x*^*2*^ =552.254; *p* < .001). The interaction between all fixed effects (CS by training by time by instrumental condition) was insignificant (*x*^*2*^ =1.524; *p* = .467) (*please see supplementary Fig*.*2*), however the significant interaction between CS by training by time (*x*^*2*^ =10.144; *p* = .006) indicates an effect of training on the change of CS-button presses (PIT effect) over time (*please see Fig*.*4*). Here, the sham training led to a decrease in button presses in negative CS trials (*estimate* = -0.361; *z* = -3.650; *p* < .001) and neutral CS trials (*estimate* = -0.211; *z* = -2.311; *p* = .021) but no significant change in positive CS trials (*estimate* = 0.001; *z* = 0.019; *p* = .985). This indicates an overall increased PIT effect after sham training. The button presses between training conditions also differed after training, here the response rate in negative CS trials was significantly higher after avoidance compared to sham training (*estimate* = 0.6304; *z* = 2.048; *p* = .041). After avoidance training, we saw a slight but insignificant increase of button presses in negative CS trials (*estimate* = 0.086; *z* = 0.967; *p* = .334). In addition, the interaction effect of time by training (*x*^*2*^ =11.290; *p* < .001) as well as time by instrumental condition (*x*^*2*^ =6.977; *p* = .008) was significant. Here, the overall button presses were significantly higher at pre training compared to post training AAT after sham training (*estimate* = 0.448; *z* = 3.360; *p* < .001) and button presses in no collect trials were significantly higher before AAT training than after training (*estimate* = 0.280; *z* = 2.641; *p* = .008).

There were no effects of training, time or an interaction on the proportion of correct trials during the forced choice task (*please see supplementary material*).

## Discussion

The goal of the current study was the assessment of an AAT modification training and its’ impact on the PIT effect in a healthy study sample. We found a significant PIT effect that did not translate into an approach or avoidance bias operationalized with a joystick AAT. Training did not result in a significant change in approach and avoidance propensities measured by AAT but exploratory analyses indicate a reduced approach towards positive CS in the avoidance training group. Our results further showed that avoidance training did not significantly reduce the PIT effect but an increased PIT effect was seen after sham training. However, avoidance training led to a significantly higher response rate towards negative CS compared to sham training.

As expected, we found robust significant PIT effects in accordance with previous reports ^12,35^. Instrumental behavior was enhanced by positive Pavlovian cues, while negative cues constrained instrumental responding. Contrary to our hypothesis, we did not see the effect of Pavlovian cue presentation translate to approach/avoidance behavior in our joystick AAT. We expected the positively valued cues to be approached faster and to be avoided slower and vice versa for negatively valued cues. The lack of significant bias towards the conditioned stimuli might be due to various reasons. For instance, we are using an irrelevant feature version, i.e. participants respond to a feature related to image orientation (horizontal/vertical) and not to the content of the image per se. Studies have made a strong case for the relevant feature version of the AAT for bias measurement ^40,41^. As a result, our implicit AAT might not capture possible bias effects as attention is not drawn to the content of the stimuli. In contrast to the transfer part of the PIT task, the attributes of the stimuli per se might not capture the subjects’ attention in the AAT. In the PIT task, the Pavlovian cues are tiled over the background while subjects are instructed to acknowledge their presence and their associated value but to respond to the instrumental stimuli only. The AAT instructions refer to the orientation of the image only but not to content - in line with previous AAT training studies ^42,43^.

In line with the above, instructions have been found to be an important predictor of approach bias and overall significance was related to individuals awareness of stimuli valences ^44^. From an associative stand, AAT bias was attributed to impulsive, automatic processes and training could alter implicit associations towards cues ^18^. This view is now challenged by a vast body of research that propose the idea of inferential processes guiding stimulus-action tendencies ^45,46^. According to this theory, these tendencies echo learned instrumental significance that translate into goal-driven behavior ^45^. In this context, the importance of instructions can be explained by conscious learning of contingencies of the stimuli and alteration of behavior according to task demands.

In addition, there is no reward for behavior in the AAT while participants receive monetary recompence for both the presentation of the Pavlovian cues as well as instrumental responding (although no direct feedback is given after the trials). Another explanation could relate to the operationalization of the response in PIT and AAT i.e. button pressing versus pushing/pulling of a joystick. Conversely, a PIT paradigm that instrumentalized joystick approach/avoidance movements instead of button presses has been established and robust PIT effects have been found ^47^. We also found a significant main effect of time that suggests a learning effect reflected by faster RT in the AAT overall. After AAT training the approach and avoidance propensities towards the CS changed: overall we saw that negative CS compared to positive CS were avoided slower and approached faster. However, this was surprisingly not driven by training condition and the sham training group showed the same effect. Again, it is not clear if an approach/avoidance bias towards the CS can be established here, the overall increase in RT indicates a strong learning curve of the task demands and in line with the above, subjects might merely focus on image orientation.

When examining the PIT after training, we did however, find a significant training effect. In this case, contrary to our hypothesis, avoidance training did not reduce the PIT effect-however, after sham training the PIT effect was enhanced. The increased PIT slope after sham training did not fall in line with our expectation. As the transfer phase in this paradigm is done in nominal extinction (i.e. without direct rewarding feedback) and the CS presentation during AAT training was not linked to reward, we did not expect the magnitude of the PIT effect to increase in either condition ^48,49^. On the other hand, we found button press response towards negative CS to be increased in the group that underwent avoidance training compared to sham training. This might indicate a possible training effect in terms of increased valence of a previously negatively conditioned cue. Since this is not reflected by RT differences between both conditions during post training AAT, we hypothesize that the AAT might not capture the CS effect on instrumental behavior as the PIT paradigm does. Again, the tasks differ in terms of stimulus presentation, instruction and operationalization of the instrumental action.

As this is a preliminary proof-of concept study, the sample size per condition is rather small and we might have not detected possible effects due to low statistical power. Since, PIT effects have been proposed to be larger in clinical populations ^14,50^, inclusion of a healthy sample might have also masked possible effects. As stated above, any possible changes in instrumental behavior might be due to expectancy effects related to instructions or task set-up, however, we have no qualitative or quantitative information by the participants and think that it would be crucial to implement post-hoc questionnaires assessing the participants’ subjective judgement on task demands and cognitive strategies. In summary, our hypotheses should be investigated in a larger and clinical sample with i.e. substance abuse or other behavioral patterns marked by conflicts of behavioral goals and value assigned to stimuli.

Taken together we found CS that elicit increased instrumental responding in a PIT task to not affect action bias in an AAT. Sham control training led to a significant increase of the PIT effect, possibly due to memory reconsolidation effects. Modification of the PIT effect was seen in those participants who were trained to approach negative CS, showed increased instrumental responding towards those cues in the PIT in contrast to sham training participants. These results indicate that implicit training can alter the strength of the influence of Pavlovian CS on instrumental behavior and might render AAT training beneficial in reducing environmental triggers of maladaptive behavior.

## Acknowledgements

This work was partially supported by the German Research Foundation (DFG, Project-ID 402170461 – TRR 265; 390688087 – EXC 2049) and the China Scholarship Council (CSC grant 201806750014).

## Author contributions

A.B. and N.R.-S. conceptualized the study. A.R. wrote the first draft of the manuscript and prepared all tables and figures. A.B., N.R.-S., K.C. and A.R. reviewed and edited the manuscript. A.R. finalized and submitted the manuscript.

## Data availability statement

The datasets generated during and/or analyzed during the current study are available from the corresponding author on reasonable request.

