## Supplementary Material for "Modifying Pavlovian-To-Instrumental Transfer By Approach Avoidance Training In Healthy Subjects – A Proof of Concept Study"

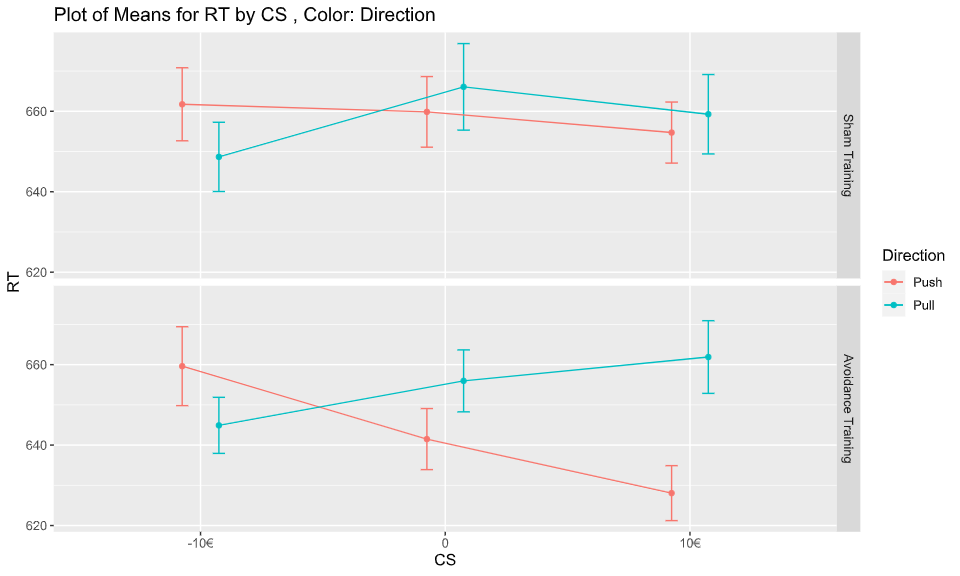

**Supplementary figure 1.** Graph depicting the effect of training on the AAT RT post training.

*Instrumental condition: Not collecting*

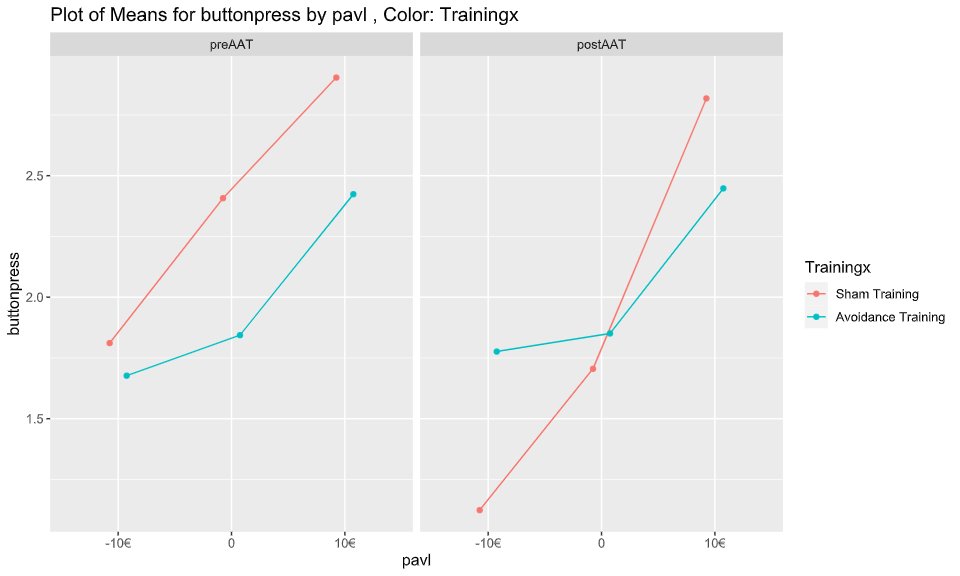

*Instrumental condition: Collecting*

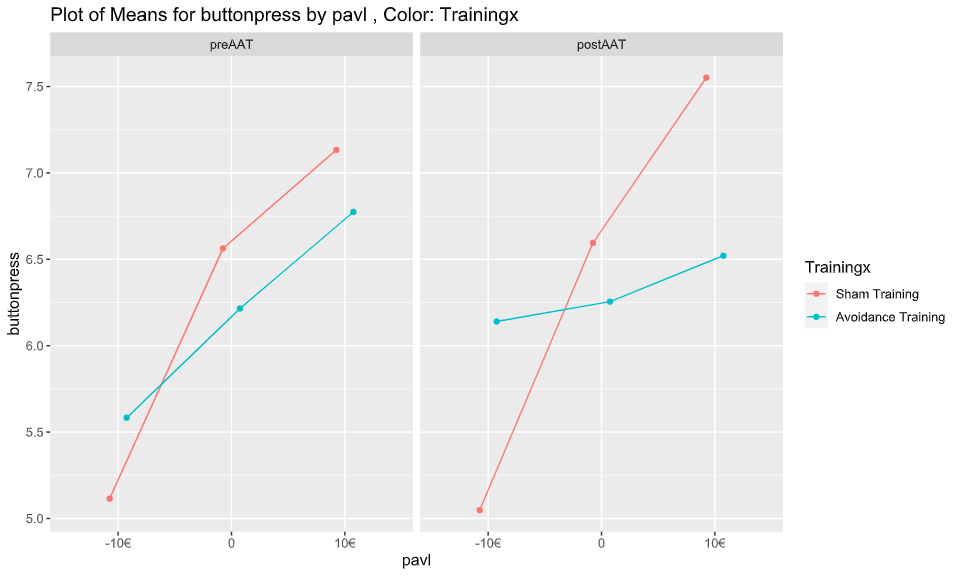

**Supplementary figure 2.** Effect of sham and avoidance training on PIT buttonpresses divided by instrumental condition (collect/not collect).

**Query trials**

To assess the effect of the training condition over time on the number of correct/incorrect choices during the query trials, we used a logistic GLMM to assess the interaction between time by training. There was no difference between pre and post AAT (*F* = 0.650; *p* = 0.421) as well as nature of training (*F* = 1.209; *p* = 0.280) on forced choice task accuracy. Additionally the interaction between time and training was insignificant as well (*F* = 1.333; *p* = 0.249), indicating no difference between the conditions over time on forced choice accuracy (*please see supplementary table 2 and supplementary figure 3*).

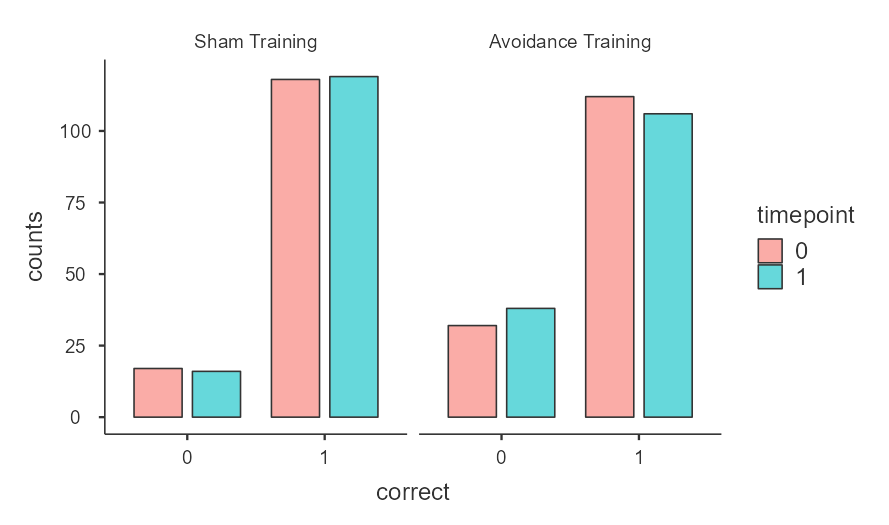

**Supplementary Figure 3.** Distribution of correct trials pre/post AAT in both sham and avoidance training conditions.

|  |  |  |  |  |
| --- | --- | --- | --- | --- |
|  | | **F** | | **p** |
| Timepoint |  | 0.650 |  | 0.421 |
| Training |  | 1.209 |  | 0.280 |
| Timepoint ✻ Training |  | 1.333 |  | 0.249 |

**Supplementary Table 1**. Results of a logistic GLMM testing the effect of training and time on the proportion of correct choices in the query trials.
